# Lysosomal swelling triggers LRRK2 activity

**DOI:** 10.64898/2025.12.09.693280

**Authors:** Tuyana Malankhanova, Zhiyong Liu, Samuel Strader, Weiping Wang, Kiwoon Sung, Zhong Li, Shawn M. Ferguson, Andrew B. West

## Abstract

LRRK2 is implicated in lysosomal functions, but the physiological upstream cues that engage endogenous LRRK2 activity are incompletely defined. Here we show that lysosomal swelling serves as a selective and reversible trigger for LRRK2-mediated Rab phosphorylation, without requiring membrane damage. Acute inhibition of PIKfyve, but not the general disruption of phosphoinositide signaling, induces the robust accumulation of phosphorylated Rabs across endolysosomal membranes. Rescue of swelling through pharmacological restoration of lysosomal ionic imbalances from PIKfyve inhibition suppresses LRRK2 activation without restoring lysosomal function. Mechanical lysosomal swelling from indigestible osmolyte uptake causes a dose-dependent increase in LRRK2-mediated Rab phosphorylation on both swollen and non-swollen lysosomes. Together, these findings identify LRRK2 as a sensor of lysosomal volume and mechanical stress, not specifically membrane damage or PIKfyve inhibition. As lysosomal swelling is a shared pathological feature across *LRRK2*-linked diseases, these results reframe LRRK2 as part of an endolysosomal surveillance system responsive to lysosomal distension.

## Introduction

Leucine-rich repeat kinase 2 (LRRK2) is a large multidomain kinase linked to multiple human diseases, including Parkinson’s disease, inflammatory bowel disease, and mycobacterium infections (*1*). With high *LRRK2* expression in many types of phagocytic cells in the innate immune system, including macrophages in the periphery and microglia in the brain, LRRK2 is thought to contribute to immunological responses through modification of vesicular trafficking and endolysosomal function (*2*). LRRK2 phosphorylation of Rab substrates regulates vesicle identity, trafficking, and recycling. Pathogenic hyperactivation of this pathway is strongly implicated in neurodegenerative, inflammatory, and infectious diseases (*3*). The expression of pathogenic LRRK2 protein with missense mutations that activate Rab phosphorylation have been linked to a loss of lysosomal homeostasis in different cell types (*4*, *5*). However, the upstream physiological cues that trigger wild-type LRRK2 activation at different endolysosomal membranes remain poorly understood. Candidate LRRK2 activity triggers recently reported include oxidative stress, membrane permeabilization, Rab12, TRPML1 activation, GABARAP and autophagy (*6*).

Lysosomes in immune cells integrate proteostatic and metabolic signals through their tightly regulated ionic and lipid environments, enabling the processing of diverse cargos that range from pathogens to misfolded and aggregated proteins. A critical unresolved question is whether LRRK2 activation of Rab phosphorylation on different types of endolysosomal vesicles requires typical forms of lysosomal dysfunction such as deacidification, catabolic failure, or membrane damage, or whether LRRK2 responds to more generalized lysosomal biophysical or mechanical changes. Phosphoinositide metabolism represents one such broad regulatory axis for lysosomal function and repair. Phosphoinositide species including PI3P, PI4P, and PI(3,5)P_2_ govern cargo sorting, membrane remodeling, ion channel regulation, and acidification (*7*). PI(3,5)P_2_, generated by the kinase PIKfyve, is enriched on late endosomes and lysosomes and is essential for maintaining vesicle volume, ion balance, and degradative capacity (*8–11*). Inhibition of PIKfyve activity alters lysosomal morphology through inducing ionic imbalance and swelling (*12–14*). In contrast, perturbations to PI3P or PI4P pools alter early endosomes, autophagy initiation, trafficking at the trans-Golgi network, and membrane repair (*7*).

Swollen lysosomes are pathological characteristics of disorders genetically linked or associated with the *LRRK2* gene (*15–17*). Mechanical enlargement of lysosomes can occur independently of phosphoinositide signaling through the intraluminal accumulation of indigestible osmolytes, a hallmark of chronic phagocytic burden and defective cargo catabolism (*18*, *19*). Swollen lysosomes are observed in cells challenged with pathogens, oxidized lipids, protein aggregates, or age-related debris, as well as macrophages acquiring senescent or exhausted phenotypes (*20–24*). However, it has been unclear whether lysosomal swelling itself constitutes an upstream signal sufficient to activate LRRK2-mediated Rab phosphorylation, or if swelling correlates with other initiating lysosome-derived stress signals such as membrane rupture, ionic collapse, or disrupted lipid composition.

Here, we unravel how selective perturbations in phosphatidylinositol signaling, ion balance, and mechanical load regulate the activation state of LRRK2-mediated Rab phosphorylation. Through pharmacological, genetic, and biochemical approaches that manipulate phosphoinositide signaling, osmotic lysosomal swelling, proton flux, and measures of isolated lysosomes, we show that LRRK2 responds robustly to lysosomal volume expansion in the absence of membrane permeabilization or canonical functional deficits. We further demonstrate that phosphorylated Rabs across the endolysosomal compartment accumulate without proportional increases in LRRK2 enrichment on lysosomal membranes, indicating that lysosomal membrane recruitment is not rate-limiting under these conditions. Together, our findings establish lysosomal swelling as a reversible and selective mechanical trigger for LRRK2-mediated Rab phosphorylation, independent of PI3K/PI4K pathways, luminal pH, or membrane damage signaling. This framework identifies LRRK2 as a sensor of aberrant lysosomal volumes and offers a unified model for understanding how diverse infectious, inflammatory, and protein aggregation disorders may converge on LRRK2-Rab signaling pathways.

## Results

### PIKfyve-inhibition and not general PI3K pathway disruption activate LRRK2-dependent Rab phosphorylation without inducing lysosomal damage

Phosphatidylinositol signaling pathways play a central role in the regulation of endolysosomal membrane identity, trafficking, and maturation (*25–27*). LRRK2-mediated Rab substrate phosphorylation increases with lysosomal membrane permeabilization and stress (*28*, *29*). Recent studies suggest phosphoinositide signalling is essential for rapid lysosome membrane repair (*30*). To gain insight into how phosphatidylinositol signaling may regulate LRRK2 activity, we compared the effects of several phosphoinositide kinase (PIK) inhibitors that disrupt distinct lipid pools including PI3P, PI4P, and PI(3,5)P_2_ to determine which perturbations may regulate LRRK2 activity prior to or independent of lysosomal membrane permeabilization (Fig. 1A). Primary macrophages express high levels of both LRRK2 protein and Rab10, a major LRRK2 substrate (*2*, *31*, *32*). Immunocytochemical evaluation of primary mouse macrophages revealed that PIKfyve inhibition, but not PI3K or PI4K inhibition, substantially increased Rab10 phosphorylation on threonine 73 (later referred to as pRab10) in the endolysosomal compartment (Fig. 1B-D). PIKfyve inhibition did not increase the proportion of the lysosome damage coordinator galectin-3 (Gal3) co-localized with LAMP1 (Fig. 1C), suggesting PIKfyve inhibition did not cause substantial lysosomal membrane damage under these conditions. Since phosphatidylinositol pathway imbalances are known to alter Akt-pathway signaling, for example phospho-Akt repression with wortmannin treatment (*33*), we measured phospho-Akt levels in macrophages after treatment and observed decreases with apilimod and wortmannin treatment (Fig. S1A, B). GSK-A1 treatment produced an unexpected increase in phospho-Akt levels, which may relate to timing and dose in the treated macrophages. These results suggest that the dosages and duration of compound exposure successfully engaged the phosphatidylinositol pathway, despite GSK-A1 and wortmannin having no effect on LRRK2 activation. Time-course experiments revealed that pRab10 accumulation increased progressively between 1.5 and 3 hours after PIKfyve inhibition, and that pRab10 levels returned to baseline within 3 hours after washing out apilimod (Fig. 1D; Fig. S1C). Collectively, these results indicate that PIKfyve-inhibition causes a reversible LRRK2 activation without causing overt lysosomal membrane damage. In agreement with the pRab10 immunofluorescence measures in cells, immunoblotting measures of the ratio of pRab10 to total Rab10 after PIKfyve inhibition showed marked increases, whereas wortmannin or GSK-A1 treatments produced no changes (Fig. 1E, F). Moreover, the LRRK2 kinase inhibitor PFE-360 completely abolished pRab10 without affecting total LRRK2 or total Rab10 levels, confirming that PIKfyve inhibition effects are triggering LRRK2 kinase activity. GSK-A1 or wortmannin treatments did not affect basal pRab10 levels that are LRRK2 kinase dependent (Fig. 1 E, F).

**Figure 1.**
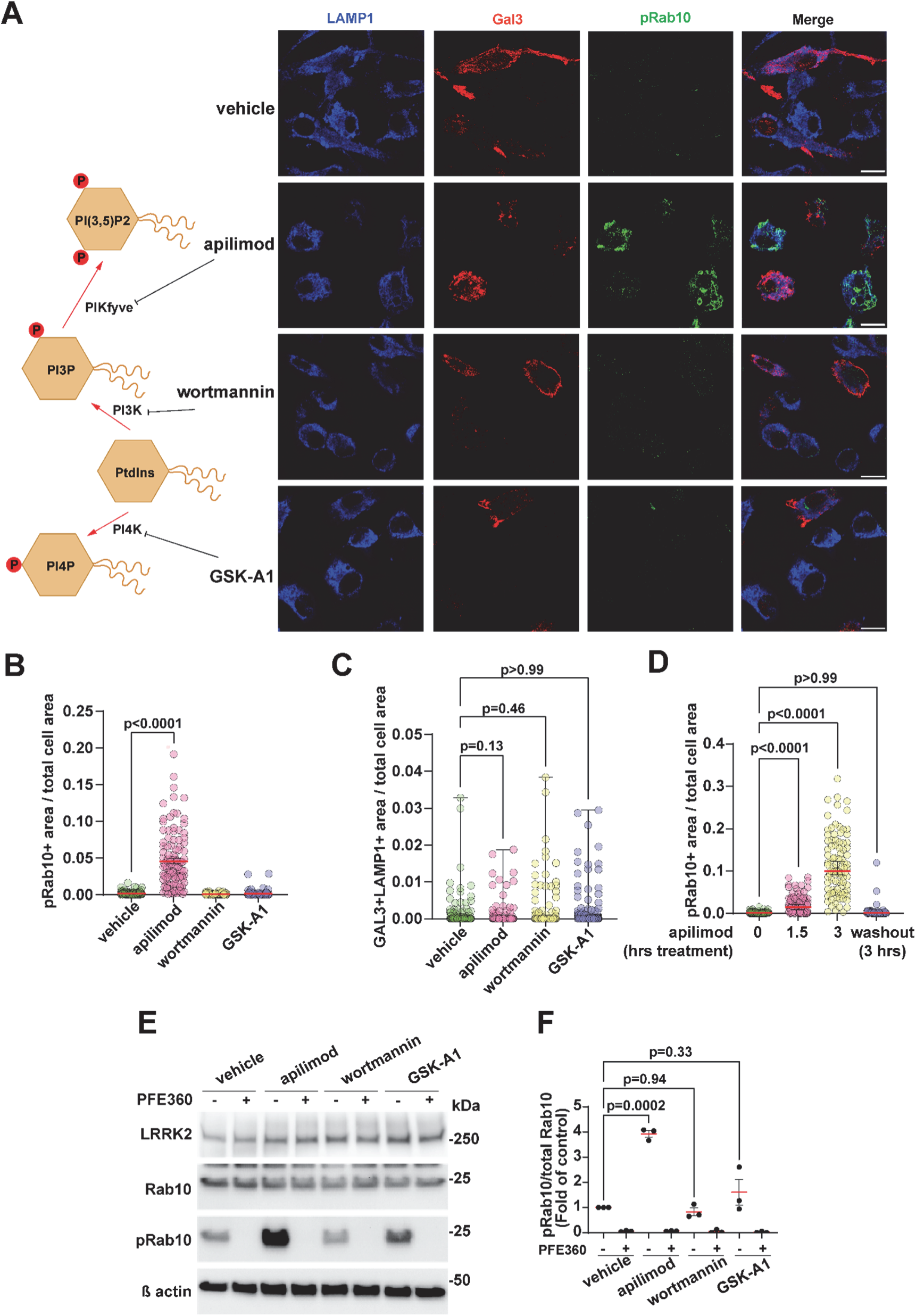
PIKfyve-specific disturbance of lysosomes rather than general disruption of the PI3K pathway activates LRRK2-dependent Rab phosphorylation without inducing lysosomal damage. **(A)** Representative confocal images of bone marrow-derived macrophages (BMDMs) immunostained for pRab10 (green), Galectin-3 (red), and LAMP1 (blue) after 2 h treatment with vehicle (DMSO, 0.01%), wortmannin (100 nM), GSK-A1 (100 nM), or apilimod (100 nM). Scale bar, 10 µm. **(B)** Quantification of pRab10⁺ area per total cell area in BMDMs showing significant increase of pRab10⁺ area in apilimod-treated cells, where each dot represents pRab10⁺ area per total cell area in one cell. **(C)** Quantification of Gal3⁺/LAMP1⁺ area per total cell area in BMDMs showing no significant change across treatments, where each dot represents Gal3⁺/LAMP1⁺ area per total cell area in one cell. **(D)** Quantification of pRab10⁺ area per total cell area in BMDMs showing pRab10 signal increased progressively with exposure time (1.5 h < 3 h) and returned to control levels after 3 h washout of 1.5 h exposure, where each dot represents pRab10⁺ area per total cell area in one cell. **(E)** Immunoblot analysis of LRRK2, pRab10, Rab10, and β-actin following 2 h treatment with vehicle (DMSO, 0.01%), wortmannin (100 nM), GSK-A1 (100 nM), or apilimod (100 nM) with or without the LRRK2 inhibitor PFE-360 (200 nM). **(F)** Quantification of pRab10/total Rab10 ratios, where each dot represents immunoblot analysis of one biological replicate. For (B), (D) and (F), data are mean ± SEM, *N* = 3 independent biological replicates. Statistical significance was assessed using one-way ANOVA with Dunnett’s post-hoc test. For (C)., data are min to max, *N* = 3 independent biological replicates, Kruskal–Wallis test. At least 30 cells per one biological replicate of immunofluorescence staining analysis were analyzed.

PIKfyve inhibition is known to elicit prominent lysosomal fission defects (*13*, *34*), and the loss of expression or activity of the Ca^2+^ release channel TRPML1 also causes marked lysosomal fission and tubulation impairment (*8*, *35*, *36*). The TRPML1 agonist ML-SA1 activates LRRK2, potentially through direct interactions with GABARAP as illustrated in Fig. 2A (*37*). To determine whether PIKfyve inhibition activates LRRK2 in a TRPML1, GABARAP, and/or Rab12-dependent mechanism, we generated knockout macrophage cells for each of these proteins (Fig. S2). ML-SA1 promotes Ca^2+^ release, TFEB activation, and lysosome tubulation (*35*, *38*). However, TRPML1 activation in ML-SA1 treatment did not rescue PIKfyve inhibition effects on LRRK2 activity. Instead, ML-SA1 caused a mild LRRK2 activation in pRab10 accumulation on its own (Fig. S2A, B). Knockout of TRPML1 expression did not affect LRRK2 activation in response to PIKfyve inhibition (Fig. 2B, C). As expected, in TRPML1 knockout macrophages, treatments with ML-SA1 had no effect on pRab10 levels, verifying the specificity of ML-SA1 on TRPML1 at drug concentrations used in this study. Knockout of either Rab12 or GABARAP (Fig. S2C-E) partially reduced the upregulation of pRab10 caused by PIKfyve inhibition (Fig. 2D-G), without significantly affecting LRRK2 expression. In all three knockout cell lines, TRPML1, GABARAP, and Rab12, homeostatic levels of pRab10 levels were similar to those measured in control macrophages (Fig. 2D-G). Together, these results suggest that PIKfyve inhibition may be activating LRRK2 and Rab phosphorylation through a mechanism distinct from lysosomal membrane damage, GABARAP, or TRPML1 function (*35*)

**Figure 2.**
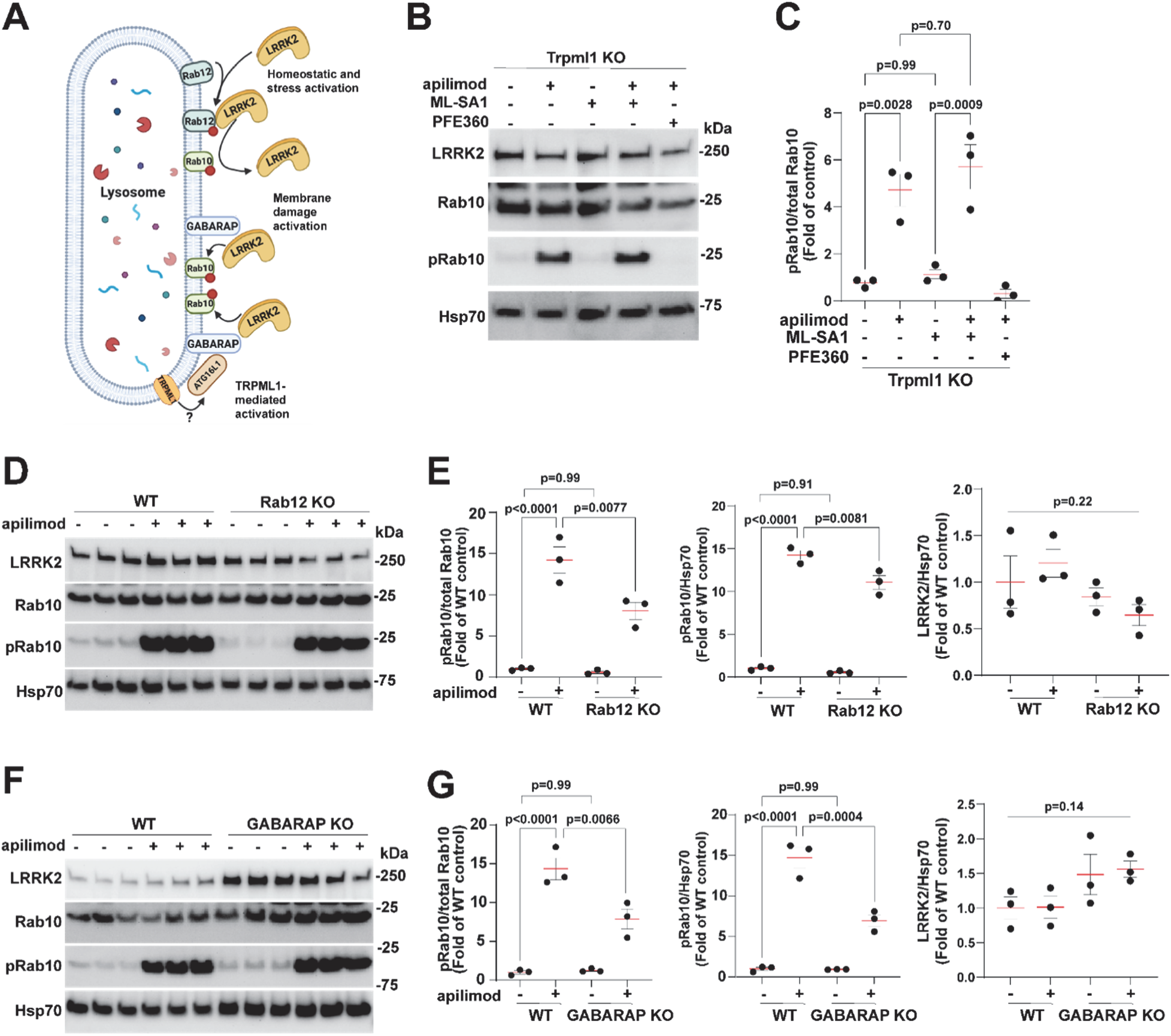
PIKfyve inhibition activates LRRK2-dependent Rab phosphorylation in a Rab12, GABARAP, and TRPML1-independent pathway. **(A)** Schematic illustration of potential lysosomal routes through which LRRK2 could phosphorylate Rab10: (i) Rab12-dependent signaling, (ii) GABARAP-linked membrane-damage activation, and (iii) TRPML1-mediated Ca²⁺ release. **(B)** Representative immunoblots of LRRK2, pRab10, Rab10, and Hsp70 in TRPML1 knockout RAW 264.7 cells following 2 h treatment with vehicle (DMSO, 0.01%), ML-SA1 (10 µM),apilimod (100 nM) and PFE-360 (200 nM) for 2 h as indicated. **(C)** Quantification of pRab10/total Rab10 ratios, where each dot represents immunoblot analysis of one biological replicate. **(D)** Representative immunoblots of LRRK2, pRab10, Rab10, and Hsp70 in wild-type (WT) and Rab12 knockout RAW 264.7 cells following 2 h treatment with vehicle (DMSO, 0.01%) or apilimod (100 nM) for 2 h. **(E)** Quantification of pRab10/total Rab10, pRab10/Hsp70, and LRRK2/Hsp70 ratios, where each dot represents immunoblot analysis of one biological replicate. **(F)** Representative immunoblots of LRRK2, pRab10, Rab10, and Hsp70 in wild-type (WT) and GABARAP knockout RAW 264.7 cells following 2 h treatment with vehicle (DMSO, 0.01%) or apilimod (100 nM) for 2 h. **(G)** Quantification of pRab10/total Rab10, pRab10/Hsp70, and LRRK2/Hsp70 ratios, where each dot represents immunoblot analysis of one biological replicate. For (C), (E), and (G), data are presented as mean ± SEM from N = 3 independent biological replicates. Statistical significance was assessed using one-way ANOVA with Tukey’s post-hoc test.

### PIKfyve inhibition depletes macrophage lysosomal bis(monoacylglycerol)phosphate (BMP) levels

Endolysosomal phosphoinositide PI(3,5)P_2_ is tightly linked to lysosomal lipid homeostasis (*39*, *40*). Bis(monoacylglycerol)phosphate (later referred to as BMP) lipids are enriched on late endosomes and lysosomes and support intraluminal vesicle formation and degradative capacity (*41*, *42*). Perturbations in lysosomal lipid homeostasis, membrane composition or maturation state alter BMP levels (*43*, *44*). LRRK2 kinase inhibition or genetic knockout leads to increased cellular BMP levels (*45–47*). To determine whether PIKfyve activity is necessary for the maintenance of normal BMP levels in macrophages, confocal imaging demonstrated a marked reduction in intracellular BMP after PIKfyve inhibition (Fig 3A, B). In contrast to PIKfyve inhibition, wortmannin or GSK-A1 treatments had modest or no effects on BMP in LAMP1-positive compartments (Fig. 3C). Mass spectrometry measures of di-22:6-BMP lipids, a highly polyunsaturated BMP species prone to rapid turnover and peroxidation (*48*), confirmed depletion of BMP lipids caused by apilimod treatment in the evaluation of total cellular lysates (Fig 3D). Thus, PIKfyve inhibition leads to pRab accumulation and BMP depletion, the opposite of the effects seen with loss of LRRK2 expression or activity, which reduces pRab levels and elevates intracellular BMP (*47*).

**Figure 3.**
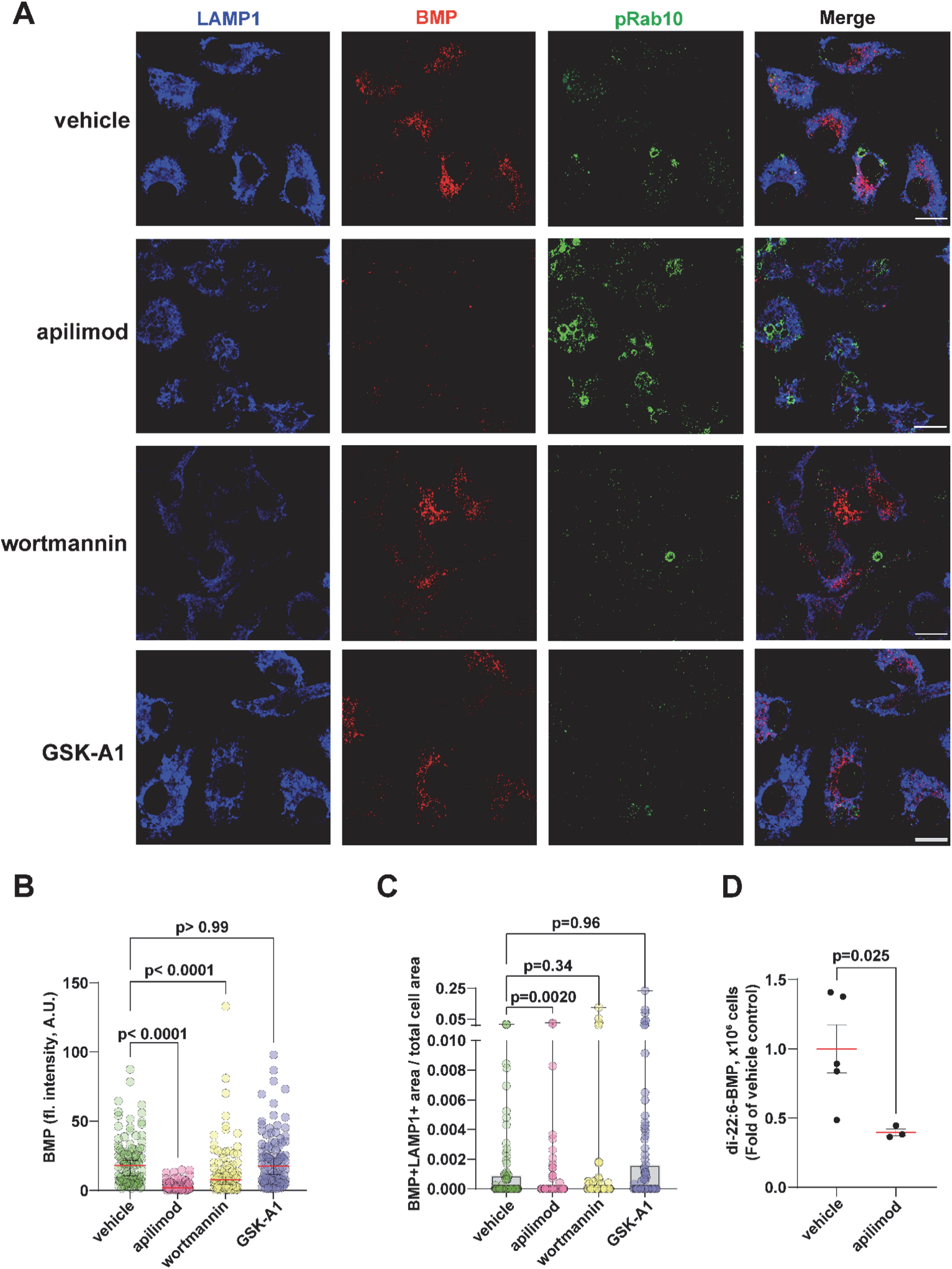
PIKfyve inhibition depletes macrophage lysosomal bis(monoacylglycerol)phosphate (BMP) levels. **(A)** Representative confocal images of BMDMs immunostained for pRab10 (green), BMP (red), and LAMP1 (blue) after 2 h treatment with vehicle (DMSO, 0.01%), wortmannin (100 nM), GSK-A1 (100 nM), or apilimod (100 nM). Scale bar, 10 µm. **(B)** Quantification of BMP signal intensity per cell in BMDMs showing significant decrease of BMP intensity in wortmannin and apilimod-treated cells, where each dot represents BMP intensity in one cell. **(C)** Quantification of BMP⁺/LAMP1⁺ area per total cell area in BMDMs showing significant decrease in apilimod-treated cells, where each dot represents BMP⁺/LAMP1⁺ area per total cell area in one cell. **(D)** BMP (di-22:6-BMP) levels were measured in WT RAW 264.7 cells treated for 2 h with vehicle (DMSO, 0.01%) or apilimod (100 nM) using LC-MS/MS. For (B), data are median with 95% CI, *N* = 3 independent biological replicates, Kruskal–Wallis test. For (C), data are min to max, *N* = 3 independent biological replicates, Kruskal–Wallis test. At least 30 cells per one biological replicate of immunofluorescence staining analysis were analyzed. For (D), data are presented as mean ± SEM; *N* = 5 for vehicle and *N* = 3 for apilimod independent biological replicates. Statistical significance was determined using an unpaired *t*-test with Welch’s correction.

### Acute V-ATPase inhibition remediates lysosomal swelling and LRRK2-Rab phosphorylation caused by PIKfyve inhibition

PIKfyve inhibition severely disrupts PI(3,5)P₂-dependent lysosomal ion balance causing substantial lysosome swelling (*10*, *49*). Brightfield imaging (Fig. S3) as well as confocal imaging (Fig. 4A, B) confirmed that all (i.e., nearly 100%) of macrophages treated with apilimod develop large swollen vesicles. We next asked whether an acute ion-reset might be sufficient to restore homeostatic levels of pRab10 in the context of PIKfyve inhibition, albeit without the restoration of normal lysosomal function or BMP lipids. The compound bafilomycin potently blocks V-ATPase-mediated proton entry. Cells treated with bafilomycin alone, or co-treated with bafilomycin and apilimod, had no swollen vesicles (Fig. 4A, B, Fig. S3). Bafilomycin effectively restores homeostatic pRab10 levels in cells in the context of PIKfyve inhibition (Fig. 4C), with a partial restoration of intracellular BMP lipids (Fig. 4D). To further probe the contribution of ionic imbalance to LRRK2 activation, we treated macrophages with the K⁺/H⁺ ionophore nigericin Nigericin collapses proton gradients across lysosomal membranes and is known to induce LRRK2 activity (*29*, *50*). Nigericin strongly induced pRab10 accumulation in macrophages to a level comparable to apilimod, and this response was completely mitigated by bafilomycin treatment (Fig. S4A, B). Confocal measures of bafilomycin-mediated restoration of pRab10 levels were confirmed by immunoblot analysis of total macrophage cellular lysates (Fig. 4E, F). Collectively, these results suggest that increased LRRK2 activity and Rab phosphorylation associated with lysosomal perturbation may be tightly coupled to lysosome swelling, not specifically PIKfyve inhibition or lysosomal functionality.

**Figure 4.**
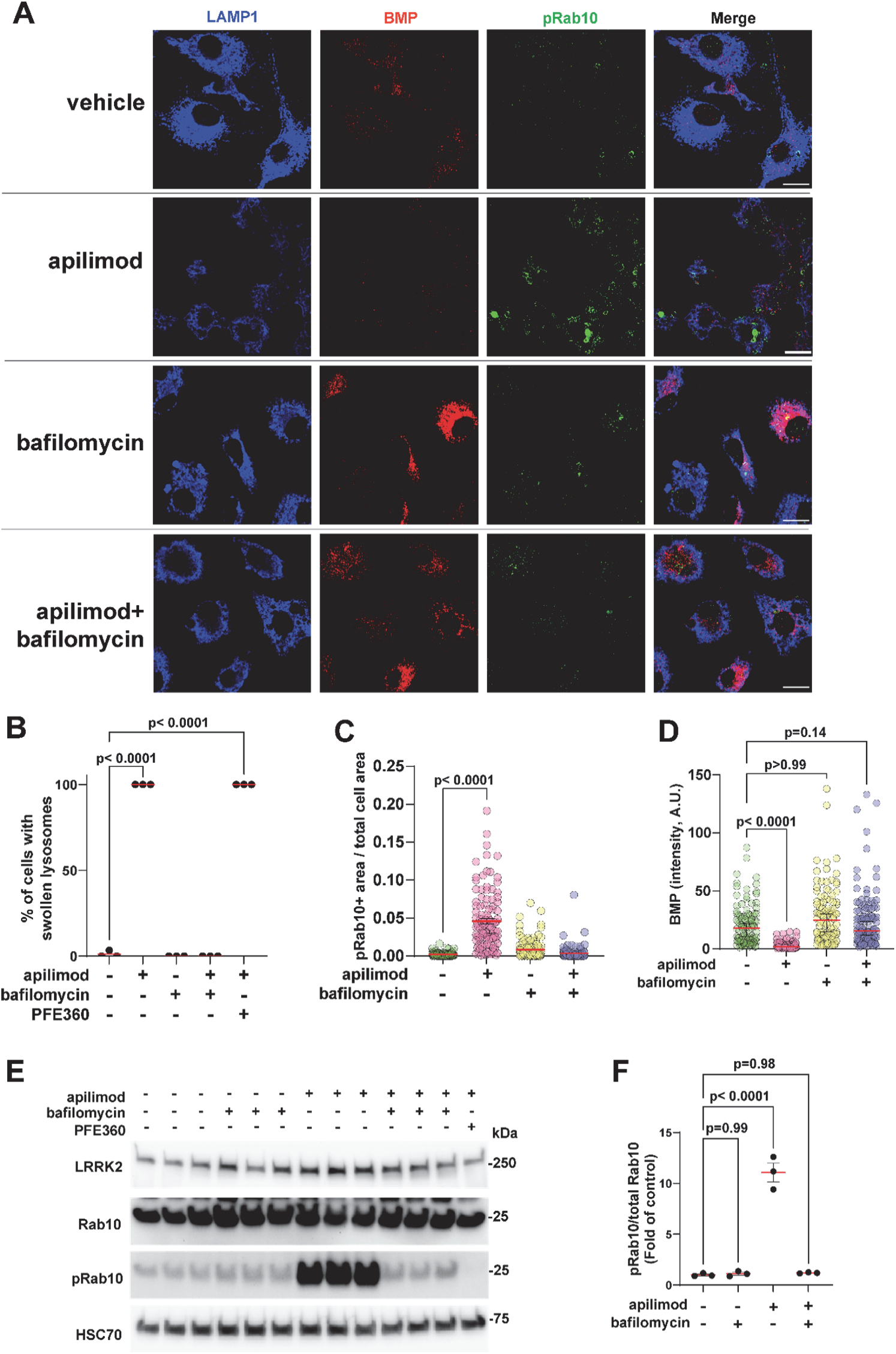
Acute V-ATPase inhibition remediates lysosomal swelling and LRRK2-Rab phosphorylation caused by PIKfyve-inhibition. **(A)** Representative confocal images of BMDMs stained for LAMP1 (blue), BMP (red), and pRab10 (green) after 2 h exposure to vehicle (DMSO, 0.01%), apilimod (100 nM), bafilomycin (200 nM), or the combination. Scale bar, 10 µm. **(B)** Quantification of the percentage of cells with swollen lysosomes (≥ 2 µm). Each dot represents the mean value from 30 cells within one biological replicate. **(C)** Quantification of pRab10⁺ area per total cell area in BMDMs showing that bafilomycin completely blocks apilimod-induced increase of pRab10⁺ area, where each dot represents pRab10⁺ area per total cell area in one cell. **(D)** Quantification of BMP signal intensity per cell in BMDMs showing that bafilomycin neutralizes apilimod-induced BMP reduction, where each dot represents BMP intensity in one cell. **(E)** Immunoblot analysis of LRRK2, Rab10, pRab10, and HSC70 in lysates from WT RAW 264.7 cells treated with apilimod (100 nM), bafilomycin (200 nM), or PFE-360 (200nM) as indicated. Each dot represents immunoblot analysis of one biological replicate. **(F)** Quantification of pRab10/Rab10 ratios shows that apilimod markedly elevates pRab10, and this effect is blocked by either bafilomycin or PFE-360. For (C), (D) and (G), data are mean ± SEM, *N* = 3 independent biological replicates, one-way ANOVA test. For (E)., data are median with 95% CI, *N* = 3 independent biological replicates, Kruskal–Wallis test. At least 30 cells per one biological replicate of immunofluorescence staining analysis were analyzed.

### Mechanical lysosomal swelling activates LRRK2

Sucrose is an indigestible osmolyte that accumulates within the endolysosomal lumen and may induce osmotic swelling of lysosomes over time in macrophages (*51*). To identify the minimum concentration of sucrose and time of exposure needed to phenocopy swollen lysosomes morphologies associated with PIKfyve inhibition, increasing concentrations of sucrose (30–100 mM for 24 h) were applied to macrophages and evaluated for pRab10 changes as well as the enlargement of LAMP1⁺ vesicles (Fig. 6A, B, and Fig. S5A). The accumulation of pRab10 as measured by immunoblot analysis is closely associated with the proportion of macrophages demonstrating swollen lysosomes caused by sucrose uptake (Fig 6B-D). Further comparative analysis of apilimod- and sucrose-induced swelling showed that both treatments produced LAMP1-positive swollen vesicles, including in TMEM192/LAMP1 double-positive compartments (Fig. S5B). As expected, the LRRK2 kinase inhibitor PFE-360 completely abolished sucrose-induced Rab10 phosphorylation (Fig. 5E). 100 mM sucrose treatment has mild effects on lysosomal deacidification, despite the substantial swelling (Fig. S5C, D). While bafilomycin treatment substantially improves lysosomal pH during PIKfyve inhibition, bafilomycin has no effect on the formation of sucrose-swollen lysosomes (Fig. S5C,D) and likewise fails to prevent LRRK2 activation and pRab10 accumulation (Fig 5E-H). Collectively, these findings uncover a dose-response relationship between the accumulation of mechanically swollen lysosomes and the induction of LRRK2-mediated Rab phosphorylation.

**Figure 5.**
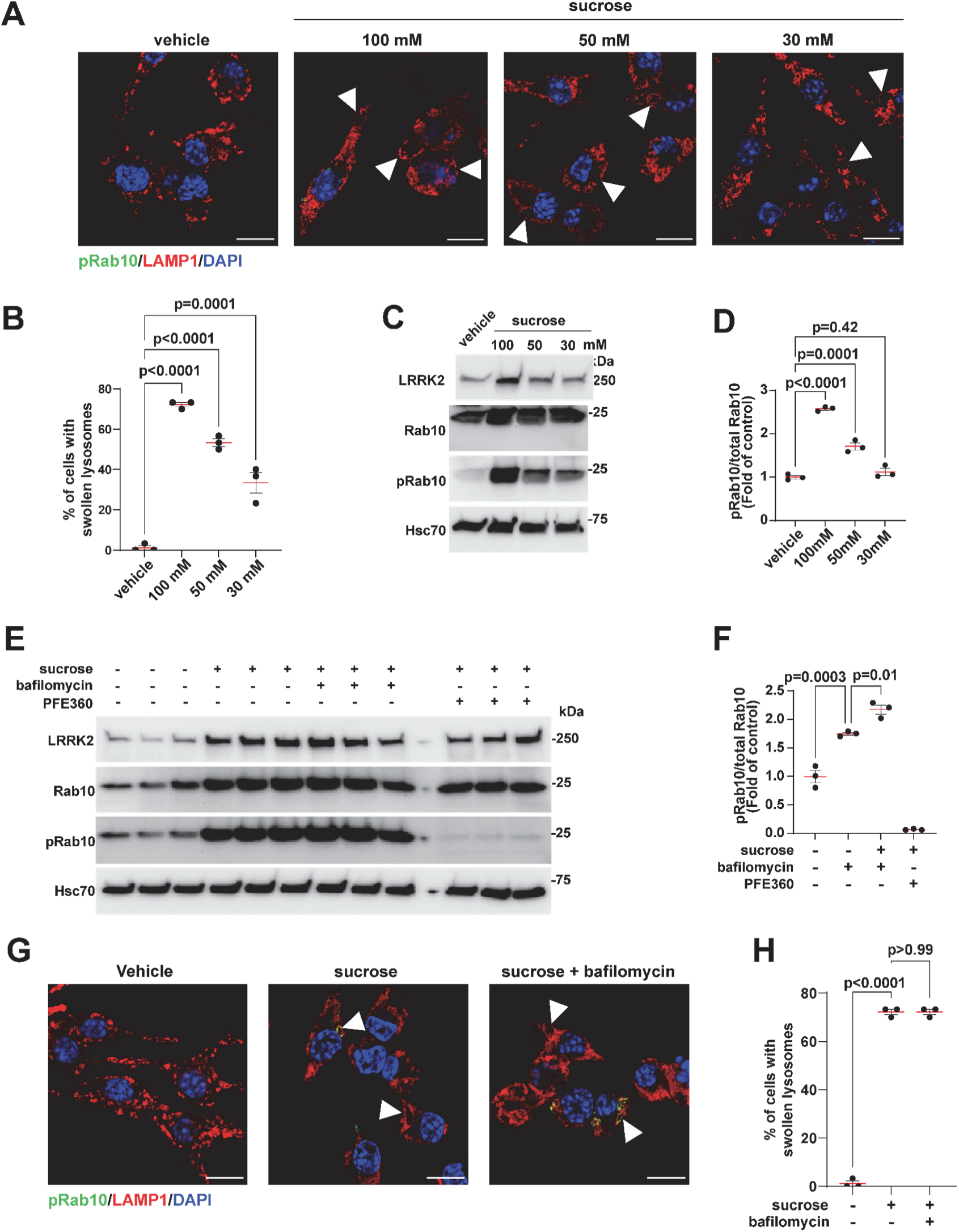
The induction of osmotic lysosomal swelling induces LRRK2–Rab phosphorylation. **(A)** Representative confocal images of RAW 264.7 macrophages stained for pRab10 (green), LAMP1 (red), and DAPI (blue) after 24 h exposure to vehicle or sucrose at 100, 50, or 30 mM. Arrowheads mark cells with enlarged LAMP1-positive lysosomes. Scale bar, 10 µm. **(B)** Quantification of the percentage of cells containing swollen lysosomes (≥ 2 µm). Each dot represents the mean from 30 cells per biological replicate. Lysosomal swelling increased with sucrose concentration. **(C)** Immunoblot analysis of LRRK2, Rab10, pRab10, and Hsc70 in lysates from WT RAW 264.7 cells treated with sucrose (100, 50, or 30 mM). **(D)** Quantification of pRab10/Rab10 ratios from cells treated with sucrose reveals significant increase of Rab10 phosphorylation. Each dot represents immunoblot analysis of one biological replicate. **(E)** Immunoblots of cells treated with sucrose (24h, 100 mM) in the presence or absence of bafilomycin (2h, 200 nM) and the LRRK2 inhibitor PFE-360 (200 nM). **(F)** Quantification of pRab10/Rab10 ratios demonstrates that PFE-360 abolished sucrose-induced pRab10 elevation, whereas bafilomycin had no effect. Each dot represents immunoblot analysis of one biological replicate. **(G)** Representative confocal images of cells with swollen lysosomes after sucrose ± bafilomycin treatment. Scale bar, 10 µm. **(H)** Quantification of the percentage of cells containing swollen lysosomes (≥ 2 µm). Each dot represents the mean from 30 cells per biological replicate. Sucrose-induced lysosomal enlargement was unaffected by bafilomycin. For (B), (D), (F) and (H), data are mean ± SEM, *N* = 3 independent biological replicates, one-way ANOVA test.

**Figure 6.**
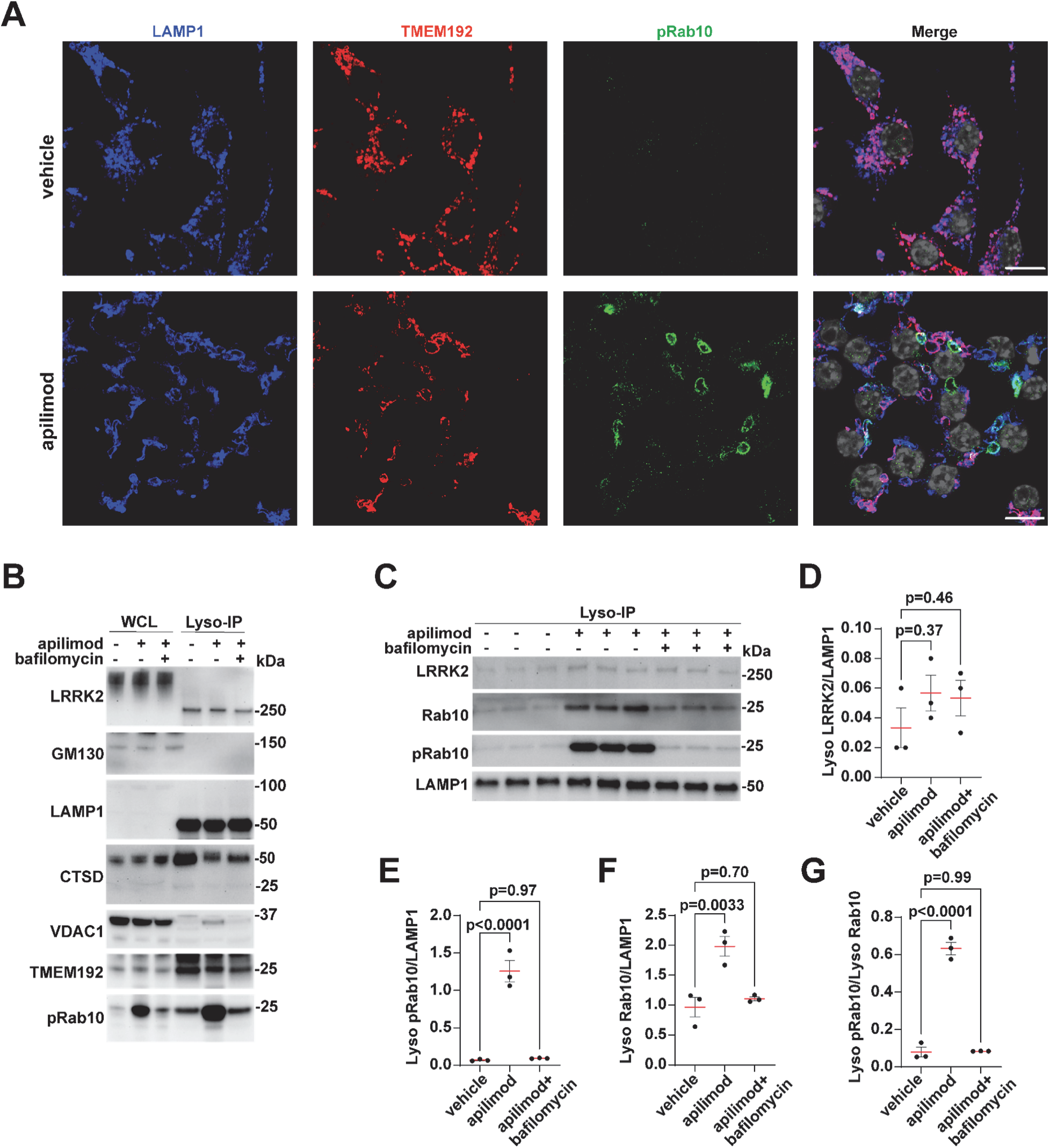
PIKfyve-inhibition does not enrich LRRK2 protein on mature TMEM192-positive lysosomal membranes. **(A)** Representative confocal images of RAW 264.7 cells stained for LAMP1 (blue), TMEM192 (red), and pRab10 (green) after 2 h treatment with vehicle (DMSO, 0.01%) or apilimod (100 nM). Scale bar, 10 µm. **(B)** Immunoblots of whole-cell lysates (WCL) and lysosome immunoprecipitates (Lyso-IP) from RAW 264.7 cells treated with apilimod ± bafilomycin. LAMP1, CTSD, TMEM192, GM130, and VDAC1 confirm fraction purity. LRRK2 is detectable in both WCL and Lyso-IP fractions but shows no significant increase at lysosomes following apilimod. **(C)** Lyso-IP immunoblots showing LRRK2, Rab10, pRab10, and LAMP1 in cells treated with apilimod and/or bafilomycin. **(D)** Quantification of lysosome-associated LRRK2 showing that LRRK2/LAMP1 ratio remains unchanged between treatments. Each dot represents immunoblot analysis of one biological replicate. **(E)** Quantification of lysosome-associated pRab10 showing that pRab10/LAMP1 ratio increases with apilimod and is abolished by bafilomycin. Each dot represents immunoblot analysis of one biological replicate. **(F)** Quantification of lysosome-associated Rab10 showing that Rab10/LAMP1 ratio reveals no significant difference. Each dot represents immunoblot analysis of one biological replicate. **(G)** Quantification of pRab10/Rab10 ratio in lysosomal fractions. For (D), (E), (F) and (G), data are mean ± SEM, *N* = 3 independent biological replicates, one-way ANOVA test.

### Lysosomal swelling does not induce a stable recruitment of LRRK2 protein to TMEM192-positive membranes

Several studies demonstrate that LRRK2 protein may be stably recruited to lysosomal membranes in response to lysosomal stress or membrane permeabilization (*52*, *53*). We and others have found that LRRK2 protein may be difficult to directly image on vesicles through confocal analysis in macrophages, for example, because of blocked epitope recognition sites in complexes on membranes, and a low proportion of total cytoplasmic LRRK2 protein associated with lysosomal membranes. The TMEM192-3xHA Lyso-IP strategy has proven effective in rapid, selective, and high-purity isolation of intact lysosomes from mammalian cells (*54*), facilitating sensitive immunoblot assessments of LRRK2 concentrations. As lysosomal swelling causes a substantial induction in LRRK2-mediated Rab phosphorylation, we investigated whether swelling might stabilize and enrich LRRK2 interactions on TMEM192-positive membranes (Fig. 6A). Evaluation of Lyso-IP isolates via immunoblot analysis demonstrated the expected accumulation of pRab10 caused by apilimod treatment, and mitigation from bafilomycin treatment (Fig 6B, C). Immunoblotting of Lyso-IP fractions for organelle-specific markers demonstrated high lysosomal purity (LAMP1, TMEM192, CTSD) and minimal contamination from Golgi (GM130) or mitochondria (VDAC1) (Fig. 6B). pRab10 accumulated extensively with LAMP1 (late endosomal and lysosomal marker) and TMEM192 (lysosomal marker). However, in contrast to pRab10, LRRK2 protein itself did not increase in abundance on TMEM192-positive membranes in response to PIKfyve inhibition (FIg 6C-G). These results suggest that stable LRRK2 recruitment to lysosomal membranes may not be the rate limiting step in LRRK2-mediated Rab phosphorylation.

## Discussion

This study reveals that the disruption of PI(3,5)P_2_, and not broader disruption of PI3K or PI4K pathways, induces a reversible and potent LRRK2 activation and Rab phosphorylation in a membrane damage-independent mechanism, consistent with recent complementary findings (*55*) showing that PI(3,5)P_2_ depletion is a selective trigger for LRRK2 signaling. In macrophages, lysosomal swelling, whether caused by reversible ionic imbalance (PIKfyve inhibition, nigericin) or osmotic loading (sucrose), drives LRRK2-mediated Rab phosphorylation. This pathway is largely uncoupled from lysosomal pH or canonical (e.g., homeostatic) lysosomal function. Lyso-IP isolation shows that lysosomal swelling causes marked pRab10 accumulations on LAMP1-positive and TMEM192-positive membranes, but without corresponding increases in TMEM192-positive membrane-associated LRRK2 protein, demonstrating that persistent physical recruitment of LRRK2 to lysosomes is not rate-limiting for the accumulation of phosphorylated Rab substrates. Collectively, these results challenge current models of LRRK2 membrane engagement and damage responses and introduce transient lysosomal swelling as a reversible LRRK2 activation pathway capable of fine-tuning Rab phosphorylation. During the final phases of writing this manuscript, it became clear that results and conclusions originally obtained at Duke University were also independently observed by researchers at Yale University as described in a recently contributed preprint (*56*). The alignment of results and conclusions independently obtained attests to the reproducibility of the findings.

Phosphatidylinositol lipids constitute a fundamental regulatory system that shapes membrane identity, coordinates vesicular trafficking, and maintains homeostasis of the endomembrane system (*7*). PI3P is a key determinant of early endosomal membrane identity, coordinating receptor sorting, vesicle maturation, and autophagy initiation (*57–60*). PI4P defines membrane identity at the trans-Golgi network and plasma membrane, regulating secretory trafficking, cargo export, and lipid exchange at membrane contact sites (*61–63*), as well as lysosomal membrane damage repair (*30*). PI(3,5)P₂ is essential for endolysosomal volume, ion homeostasis, fusion-fission dynamics, and degradation capacity (*8–11*). PI(3,5)P₂ regulates TRPML1 and other Ca^2+^ channels, as well as the CIC7 chloride channel. Through the pharmacological and genetic dissection of LRRK2 activity in macrophages presented here, we identify dysregulation of PI(3,5)P₂ as a critical factor linking LRRK2-mediated Rab phosphorylation to endolysosomal volume. Indeed, increased LRRK2 activity initiated by the formation of sucrosomes in macrophages couples Rab phosphorylation to lysosome swelling in a tightly controlled dose-dependent manner, not specifically dependent on PIKfyve inhibition or lysosomal functionality.

BMP lipids (bis(monoacylglycero)phosphate, also called LBPA) are essential for multiple core lysosomal functions, providing inner lysosomal membrane geometries and functionality essential for the prevention of lysosomal swelling (*64*). Apilimod treatment decreases BMP levels, whereas genetic knockout of LRRK2, or small molecule LRRK2 inhibitors, have a substantial effect on increasing intracellular BMP levels and diminishing phosphorylated Rab levels (*47*). However, the effects of pathogenic LRRK2 mutations on BMP lipid levels is not clear, with some reports indicating increases in intracellular BMPs associated with mutated LRRK2 expression (*65*), and some reports indicating both increases and decreases on intracellular BMP depending on the cell type and method of BMP detection (*47*). The effects of PIKfyve inhibition on lysosomal acidification may be both time and cell-type dependent. Although one past study reported mild lysosomal hyperacidification in U2OS cells following apilimod treatment (*66*), two other studies found no significant pH change in either U2OS cells or macrophages (*67*, *68*). The PIKfyve inhibitor WX8 has been described to cause lysosomal deacidification (*68*). Earlier work using another PIKfyve inhibitor YM201636 in NIH3T3 cells showed poorly acidified swollen vesicles after treatment, although the specific acidification state was reported to vary depending on the timing of fusion of swollen vesicles with acidified vesicles that existed prior to treatment (*69*). Thus, across published datasets and those presented herein, lysosomal pH outcomes with PIKfyve inhibition may vary depending on cell type and the timing of observation. If mechanical lysosomal swelling is the primary trigger for LRRK2-mediated Rab phosphorylation, as we suppose, and most pRab-positive vesicles are not themselves swollen but represent an endolysosomal network response, then BMP concentrations and lysosomal function may be largely decoupled (e.g., not well-correlated) with LRRK2-activity and pRab levels. Within this framework, otherwise conflicting results in the literature regarding LRRK2 activity, BMP levels, and lysosomal acidification, might be reconciled.

The *LRRK2* gene is genetically linked to infectious diseases, chronic inflammation, as well as neurodegenerative diseases (*32*, *70–72*). Swollen lysosomes that may trigger LRRK2-mediated Rab phosphorylation present in these diseases due to accumulation of infectious agents in lysosome compartments (e.g., mycobacteria or other pathogens), the presence of indigestible proteins including aggregates associated with neurodegenerative diseases, as well as chronic inflammation in exhausted or senescent phagocytic cells with higher levels of damaged or oxidized lipids. However, studies show that Rab phosphorylation in macrophages is not confined exclusively to mature endolysosomes of lysosomes (*2*), but occurs in early endosomes recently formed from the plasma membrane (*73*). LRRK2-mediated phosphorylation of Rab proteins may block vesicle recycling through blocking effector interactions including EHBP1L1 essential for membrane tubulation and cargo clearance, with EHBP1L1 especially important in high endocytic-load conditions (*53*, *74*, *75*). In the context of lysosome swelling and upstream Rab phosphorylation, the sequestration of deleterious cargo in LRRK2-mediated Rab phosphorylated endosomes may temporarily protect swollen lysosomes from additional fusion events that would further impair lysosome function. However, if the swollen lysosomes are not resolved, impaired endolysosome maturation and altered motility may exacerbate the handling of pathological cargo that may include pathogens or protein aggregates. Further, excessive Rab phosphorylation such as that caused by pathogenic *LRRK2* mutations may impair timely cargo delivery for degradation, signaling associated with lysosome swelling and impairment when none may exist.

LRRK2 specifically phosphorylates and interacts with activated, GTP and membrane bound, Rab proteins (*76*, *77*). A caveat to this study is that the exact mechanism that results in increased LRRK2-mediated Rab phosphorylation on vesicles after lysosomal swelling has not been identified. Rab12 has been identified as a key LRRK2 interacting protein for Rab10 phosphorylation (*78*, *79*). Following PIKfyve inhibition, Rab12 knockout macrophages lower pRab10 levels compared to control cells but still show a significant response to this stimulus. We hypothesize that the activation of other LRRK2 Rab interactors, for example Rab32 or Rab29, also linked to inflammation and neurodegenerative diseases (*80–83*), may be important to maintain substantial LRRK2 activity in Rab12 knockout cells. In consideration of results presented in a recently contributed preprint, LRRK2-Rab interactions are likely critical in the LRRK2 lysosome swelling pathway (*56*). At least in tissues affected by protein aggregation in tau and α-synuclein-linked diseases also genetically linked to *LRRK2*, increased pRab10 and pRab12 primarily co-localize with endolysosomal markers of dysfunction in the analysis of formalin-fixed embedded human brain tissues (*84*). Future studies will help clarify the role Rab phosphorylation plays in response to physiological challenges important in LRRK2-linked diseases like lysosomal swelling.

## Materials and methods

### Cell cultures

Mouse bone marrow-derived macrophages (BMDMs) were cultured as previously described (*85*). Briefly, BMDMs were generated by culturing the mouse bone marrow cells collected from 3- to 5-month-old FLAG-WT-mLRRK2 BAC (B6.Cg-Tg(Lrrk2)6Yue/J, The Jackson Laboratory) into DMEM (Gibco, Cat#11995073) supplemented with 10% fetal bovine serum (Atlanta Biological, Cat# S11150H (Q-39042)), 1x penicillin-streptomycin (Gibco, Cat#15140122), and 100 ng/mL mouse M-CSF (Biolegend, Cat#576406). RAW 264.7 (ATCC, Cat# TIB-71) and HEK293FT (Thermo Fisher, Cat# R70007) cells were cultured in DMEM supplemented with 10% fetal bovine serum and 1x penicillin-streptomycin.

### Pharmacologic inhibition

In all experiments, cells were exposed to the indicated compounds such as apilimod (100 nM, MCE, Cat# HY-14644), wortmannin (100 nM, MCE, Cat# HY-10197), GSK-A1 (100 nM, MCE, cat# HY-125118), bafilomycin (200 nM, cell Signaling Technology, Cat# 54645), nigericin (10 µM, MCE, Cat# HY-100381), ML-SA1 (10 µM, MCE, Cat# HY-108462), PFE-360 (200 nM) for 2 h, except where alternative incubation times are specified. RAW 264.7 cells were incubated with sucrose (100 mM, 50 mM, 30 mM) for 24 h.

### LC-MS/MS analysis of lipid di-22:6-BMP

Cell pellets were lysed with 300 μL 50% methanol by ultrasonic disruption (Q55 Qsonica Sonicator) for 2 x 30 seconds (in ice bath). The resulting supernatant extracts were transferred to a new 1.5-mL vial and mixed with 100 μL 100 ng/mL di-14:0-BMP, and 750 μL chloroform/methanol (3/1, v/v) with 62.5 μg/mL butylated hydroxytoluene, followed by the addition of 150 μL water (with 0.1% formic acid). After vortexing and centrifugation, the bottom layer of liquid was transferred into the well of the 1 mL 96-well plate, followed by complete dryness with nitrogen gas flow at room temperature. Subsequently, 100 μL solution (methanol with 5mM ammonia formate) was added into the plate wells to reconstitute the samples. Samples were analyzed with the Waters LC-MS instrument (Milford, MA) which is composed of the TQ-S MS system and Acquity UPLC. Software MassLynx 4.2 was used for data acquisition. The LC separation was performed on a Waters Acquity UPLC BEH C18 column (2.1 x 100mm, 1.7 μm) with mobile phase A (0.1% formic acid and 5mM ammonia formate in water) and mobile phase B (0.1% formic acid and 5mM ammonia formate in methanol/acetonitrile, 7/3, v/v). The flow rate was 0.45 mL/min. The linear gradient was as follows: 0-2min, 35%B; 3-4min, 70%B; 4.5-5min, 90%B; 5.5-7.2, 100%B; 7.5-8.8min, 100%B (0.85 mL/min); 8.95-10min, 35%B. The autosampler was set at 5°C and the column was kept at 55°C. The injection volume was 5 μL. Mass spectra were acquired with electrospray ionization in positive mode. Multiple reaction monitoring was used to measure the target signals: di-BMP 22:6 with m/z 867.1⤑ m/z 385.1; di-BMP 14:0 with m/z 667.2⤑ m/z 285.1. Software Skyline (version 23.1.0.455. www.skyline.ms) was used to process the instrument acquired data. Due to the lack of authentic di-22:6-BMP standard, the signal of di-14:0-BMP was used to quantify the level of di-BMP22:6 (*86*, *87*).

### Generation of knockout RAW 264.7 cells

*Rab12* KO and *Trpml1* KO RAW 264.7 cells were created using CRISPR/Cas9 expressing plasmids (Table S1). 1⨯10^6 cells were transfected with the plasmids using Neon Transfection System (Thermo Fisher, Cat# MPK10025) following the manufacturer’s protocol. After 24 h post-transfection GFP-positive cells were sorted and plated in 96-well plate 1 cell per well. After cell clonal expansion knockout clones were identified by immunoblotting (Rab12 KO) or Sanger sequencing (Trpml1 KO). GABARAP KO RAW 264.7 cells were previously reported (37).

### *Trpml1* KO validation

Genomic DNA was extracted using the Zymoclean Quick DNA Miniprep (Zymo Research, Cat# 3024) and amplified with LinkTaq HS Taq Polymerase (Biolink Laboratories, Cat# 16-1300MG-400). PCR products were purified using the DNA Clean & Concentrator-5 kit (Zymo Research, Cat# D4004) and submitted to Eton Bioscience for Sanger sequencing. Knockout validation was performed by analyzing sequencing traces with ICE Analysis (Synthego).

### Immunoblotting

Immunoblotting was performed as previously described (*85*). Protein lysates were analyzed using SDS–PAGE (Bio-rad, Cat# 4568096) followed by transferring to PVDF membranes (Fisher Scientific, Cat# IPFL00010) for immunoblotting with the indicated primary antibodies and HRP-conjugated secondary antibodies. Signals were developed with Crescendo ECL reagent (Millipore, Cat# WBLUR0500) on a Chemidoc MP platform (BioRad). Saturated signals on immunoblots were not detected in any experiment used for analysis (ImageLab 6.1), and representative signals used for analysis are shown in the figures. A complete list of all antibodies used in this study is provided in Table S2.

### Immunofluorescence

Immunofluorescence staining was performed as previously described (*85*). Briefly, cells were fixed with 4% paraformaldehyde (VWR, Cat# 15714-S), washed three times with PBS (VWR, Cat# 21040CM), permeabilized with 0.1% saponin (Sigma-Aldrich, Cat #S4521-25G), and blocked with 3% normal donkey serum (Abcam, Cat# ab138579-50mL) and immunostained with the primary and then secondary antibodies listed in the Table S2. DAPI dye (Thermo Fisher, Cat# D1306) was used for nuclear staining. Cell culture coverslips were mounted onto glass slides with ProLong Gold antifade reagent (Invitrogen, Cat# P36934).

### Microscopy

Immunofluorescence staining imaging was performed as previously described (*85*). Briefly, microscopy images were obtained using Airyscan on a Zeiss 880 Inverted confocal microscope with a 63× oil immersion objective. Airyscan images were processed using Zen Black 3.0 software (Carl Zeiss). Brightfield imaging was performed using Olympus VS200 Research Slide Scanner with a 60× oil immersion objective. Images were processed using Olyvia and QuPath software.

### LysoSensor Green staining

Cells were incubated with LysoSensor Green DND-189 (Thermo Fisher Scientific, Cat# L7535) at 1 µM in complete medium for 30 min at 37 °C, washed with pre-warmed PBS, and immediately imaged live on a confocal microscope. Hoechst dye (BD Biosciences, Cat# 561908) was used for nuclear staining.

### Image analysis

Fluorescence images were analyzed using a standardized CellProfiler (v4.2.8) workflow. After *RescaleIntensity*, nuclei were segmented from the DAPI channel with *IdentifyPrimaryObjects*. Whole-cell and cytoplasmic regions were generated using *IdentifySecondaryObjects* and *IdentifyTertiaryObjects*. Protein-specific puncta were detected with *IdentifyPrimaryObjects* using adaptive Otsu thresholding and declumping. All markers were quantified within their defined compartments, with pRab10 restricted to the cytoplasmic region to exclude nuclear non-specific signals. Measurements included puncta count and total positive area. Colocalization was assessed with *RelateObjects* by calculating overlapping puncta area normalized to cell area. Outputs were exported as .csv files for downstream analysis, and identical pipeline parameters were applied to all images to ensure reproducibility. For LysoSensor Assay and BMP signal intensity, fluorescence intensity was quantified from confocal images using Fiji v2.16.0. Brightfield images were analyzed by manual inspection, and cells exhibiting vesicular structures greater than 2 µm in diameter were scored as swelling-positive.

### Lentivirus production and transduction

Lentiviral particles encoding TMEM192-3xHA were produced in 293FT cells using FuGENE 6 (Promega, Cat# E2691) transfection reagent. One day prior to transfection, 4–6 × 10⁶ 293FT cells were seeded into T75 flasks to obtain ∼70–80% confluency at the time of transfection. For each T75 flask, plasmid DNA was prepared at the following amounts: 10 µg TMEM192-3xHA transfer vector (plasmid #102930, Addgene), 7.5 µg psPAX2 packaging plasmid (plasmid #12260, Addgene), and 2.5 µg pCMV-VSV-G envelope plasmid (plasmid #8454, Addgene) (20 µg total DNA). FuGENE 6 was used at a 3:1 reagent-to-DNA ratio (60 µL FuGENE 6 per 20 µg DNA). FuGENE reagent was diluted in 1 mL serum-free DMEM and incubated for 5 min at room temperature. The plasmid DNA mixture was then added to the diluted FuGENE, gently mixed, and incubated for 15–20 min to allow complex formation. The complete DNA–FuGENE complex was added dropwise to the cells and evenly distributed by gentle swirling. Cells were maintained at 37 °C and 5% CO₂. Medium was replaced after 16 h. Viral supernatants were collected at 48 h post-transfection. The harvested medium was centrifuged at 800 × g for 10 min to remove cellular debris, and the clarified supernatant was transferred to clean 50-mL tubes. Viral particles were concentrated by adding 4×Lentivirus Concentrator Solution (40% w/v PEG-8000, 1.2 M NaCl in 2×PBS (pH7.4)), at a ratio of 1 volume concentrator to 3 volumes supernatant. The mixture was shaken for 60 s and incubated with gentle rocking (∼60 rpm) overnight at 4 °C. Viral particles were pelleted by centrifugation at 1,600 × g for 60 min at 4 °C. Supernatants were carefully removed, and viral pellets were resuspended thoroughly in 1 mL of serum-free DMEM. Concentrated lentivirus was aliquoted and stored at −80 °C; repeated freeze–thaw cycles were avoided.

### Lentiviral transduction of RAW 264.7 cells

RAW 264.7 cells were seeded one day prior to transduction to reach approximately 70–80% confluency at the time of infection. Lentiviral supernatant was added to the culture medium at a 1:10 ratio and cells were incubated with virus for 6 h. 24 h following transduction, cells were selected with puromycin at 4 µg/mL for 7 days. Surviving resistant cells were subsequently maintained in a growth medium containing 4 µg/mL puromycin to preserve transgene expression. Successful transduction was verified by anti-HA immunofluorescence staining and by immunoblotting.

### Lysosomal immunoprecipitation

Lysosomal purification was performed according to the STAR Protocol by *Clément Maghe and Julie Gavard* (2024) without modifications (*88*).

### Statistical analysis

The indicated *n* refers to the number of independent preparations (biological replicates) used for immunoblotting and immunofluorescence analyses. All statistical analyses were performed using GraphPad Prism 10. Data distributions were assessed using Shapiro–Wilk tests. Statistical significance was determined using one-way ANOVA with Tukey’s post hoc comparison or, when assumptions for parametric testing were not met, Kruskal–Wallis tests with Dunn’s post hoc comparisons. *P* values < 0.05 were considered statistically significant. Specific details of statistical tests are provided in the figure legends. No blinding was performed.

## Acknowledgments

We thank Devin Clegg for contribution of GABARAP knockout macrophages, and Hilary Chester for plasmids encoding TMEM192-3xHA and lentiviral packaging plasmids.

## Data Availability

Raw data associated with the figure panels and a key resources table (KRT) is available online through the Zenodo data repository doi.org/10.5281/zenodo.17782145

---

**Table S1.**
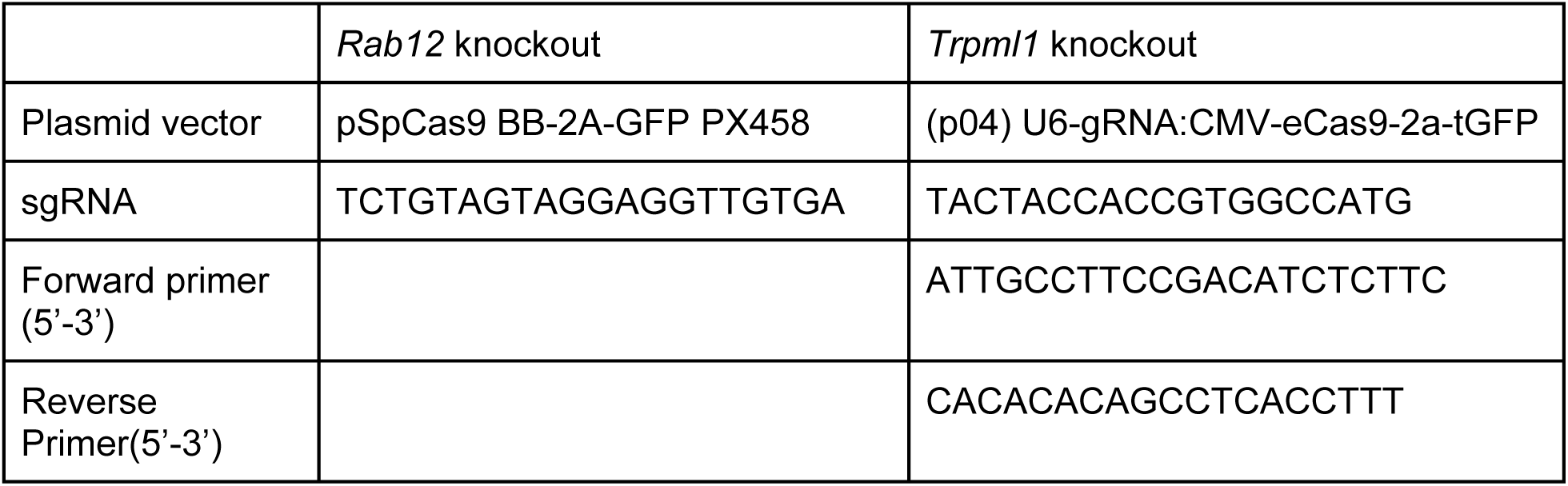
Genetic knockout reagents.

**Table S2.** Antibodies.

|  | Catalog # | Manufacturer | Dilution |
| --- | --- | --- | --- |
| anti-LRRK2 | ab133474 | Abcam | 1:2000 |
| anti-phospho-T73-Rab10 | ab230261 | Abcam | 1:1000 |
| anti-phospho-T73-Rab10 | ab241060 | Abcam | 1:1000 |
| anti-Rab10 | ab237703 | Abcam | 1:2000 |
| anti-galectin 3 | MA1940 | Invitrogen | 1:1000 |
| anti-LBPA | MABT837 | Sigma | 1:200 |
| anti-β-actin | sc-47778 HRP | Santa-Cruz | 1:5000 |
| anti-Hsp70 | PA5-28003 | Thermo Fisher | 1:3000 |
| anti-Hsc70 | sc-7298 | Santa-Cruz | 1:5000 |
| anti-GM130 | 12480 | Cell Signalling | 1:1000 |
| anti-HA | H3663 | Sigma | 1:1000 |
| anti-LAMP1 | sc-20011 | Santa-Cruz | 1:1000 |
| anti-LAMP1 | sc-19992 | Santa-Cruz | 1:1000 |
| anti-CTSD | AF1029 | Biotechne | 1:200 |
| anti-VDAC1 | ab186321 | Abcam | 1:1000 |
| anti-TMEM192 | ab185545 | Abcam | 1:2000 |
| anti-pan-Akt | 4691 | Cell Signalling | 1:1000 |
| anti-pAkt | 4060 | Cell Signalling | 1:2000 |
| anti-GAPDH | SC-32233 | Santa Cruz | 1:1000 |
| anti-Rab12 | 18843-1-AP | Proteintech | 1:1000 |
| anti-phospho Rab12 | ab256487 | Abcam | 1:1000 |
| anti-GABARAP | 13733T | Cell Signalling | 1:1000 |
| Peroxidase AffiniPure donkey Anti-Rabbit | 711-035-152 | Jackson ImmunoResearch | 1:10000 |
| Peroxidase AffiniPure Goat Anti-Mouse | 115-035-003 | Jackson ImmunoResearch | 1:10000 |
| Donkey anti-Mouse IgG (H+L) Highly Cross-Adsorbed Secondary Antibody, Alexa Fluor 647 | A31571 | Thermo Fisher | 1:1000 |
| Donkey anti-Rabbit IgG (H+L) Highly Cross-Adsorbed Secondary Antibody, Alexa Fluor™ 488 | A21206 | Thermo Fisher | 1:1000 |

## SUPPLEMENTAL FIGURES

**Figure S1.**
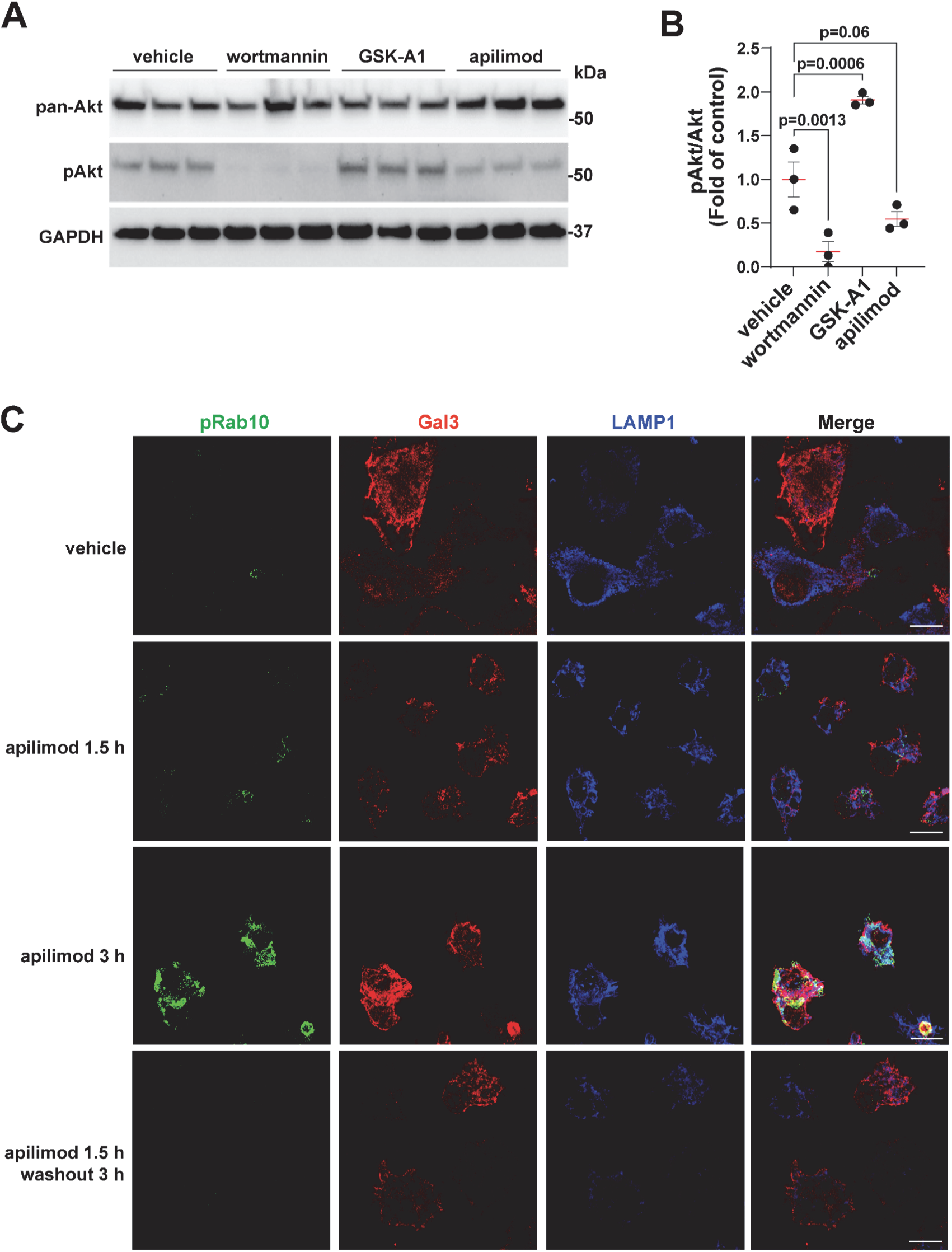
PIK-kinase inhibitor specificity and reversibility of apilimod-induced pRab10 activation. **(A)** Immunoblots of RAW 264.7 macrophages treated for 2 h with the indicated PI-kinase inhibitors: wortmannin (pan PI3K), GSK-A1 (PI4K), apilimod (PIKfyve). Blots were probed for phospho-Akt (pAkt), total Akt, and GAPDH as a loading control. **(B)** Quantification of pAkt/Akt ratios confirms that the inhibitor concentrations used were effective in modulating their intended target. Each dot represents immunoblot analysis of one biological replicate. **(C)** Representative confocal images of cells stained for pRab10 (green), Galectin-3 (red), and LAMP1 (blue) following apilimod treatment (1.5 h or 3 h) or 1.5 h treatment followed by 3 h washout. Apilimod induced pRab10 accumulation and modest Gal3 signal at 3 h, both of which returned to baseline after washout. Scale bar, 10 µm. For (B), data are mean ± SEM, *N* = 3 independent biological replicates, one-way ANOVA test.

**Figure S2.**
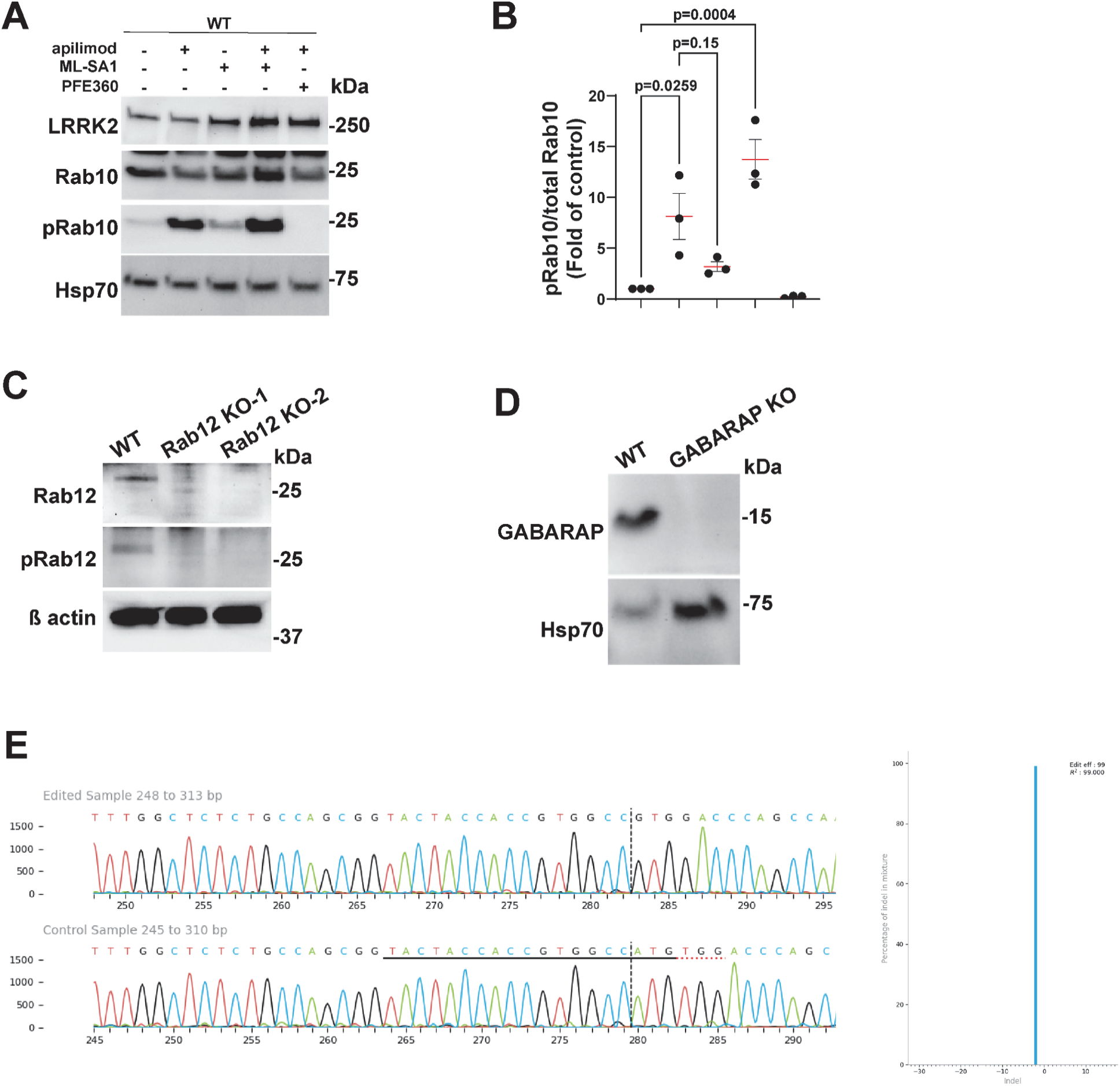
Knockout validation. **(A)** Representative immunoblots of LRRK2, pRab10, Rab10, and Hsp70 in WT RAW 264.7 cells following 2 h treatment with vehicle (DMSO, 0.01%), ML-SA1 (10 µM), apilimod (100 nM) and PFE-360 (200 nM) for 2 h as indicated. **(B)** Quantification of pRab10/total Rab10 ratios, where each dot represents immunoblot analysis of one biological replicate. Each dot represents immunoblot analysis of one biological replicate. Data are presented as mean ± SEM from N = 3 independent biological replicates. Statistical significance was assessed using one-way ANOVA with Tukey’s post-hoc test. **(C)** Immunoblots of WT and Rab12 KO RAW 264.7 macrophages. Blots were probed for pRab12, Rab12, and ß actin as a loading control. **(D)** Immunoblots of WT and GABARAP KO RAW 264.7 macrophages. Blots were probed for GABARAP, and Hcp70 as a loading control. **(E)** Knockout efficiency was quantified by ICE analysis (Synthego). ICE deconvolution indicated a KO score of 99% with 2 bp frameshift deletion.

**Figure S3.**
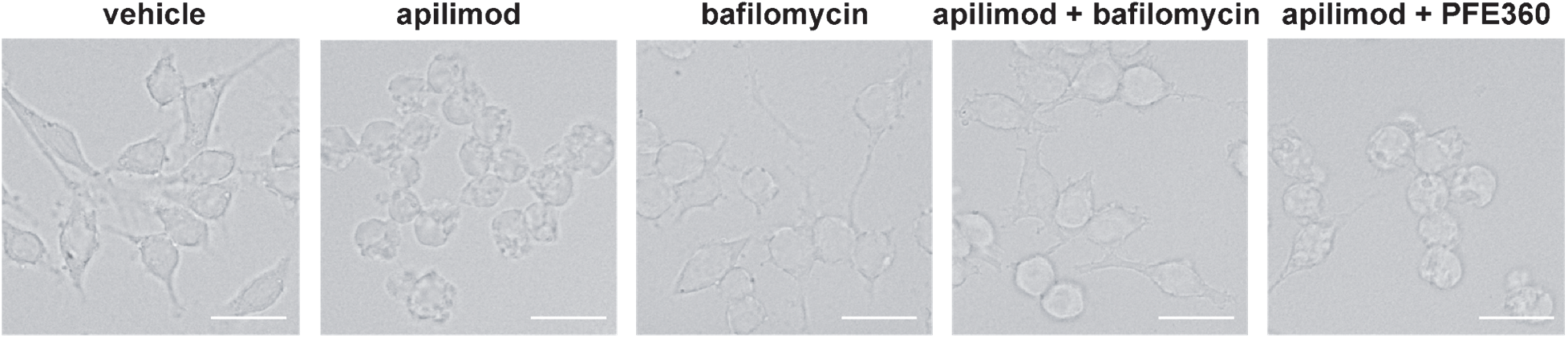
**Apilimod induces vesicle swelling that is prevented by V-ATPase inhibition** Representative brightfield images of RAW 264.7 cells after 2 h exposure to vehicle (DMSO, 0.01%), apilimod (100 nM), bafilomycin (200 nM), and the combination apilimod+bafilomycin or apilimod+PFE-360 (200 nM). Scale bar, 20 µm.

**Figure S4.**
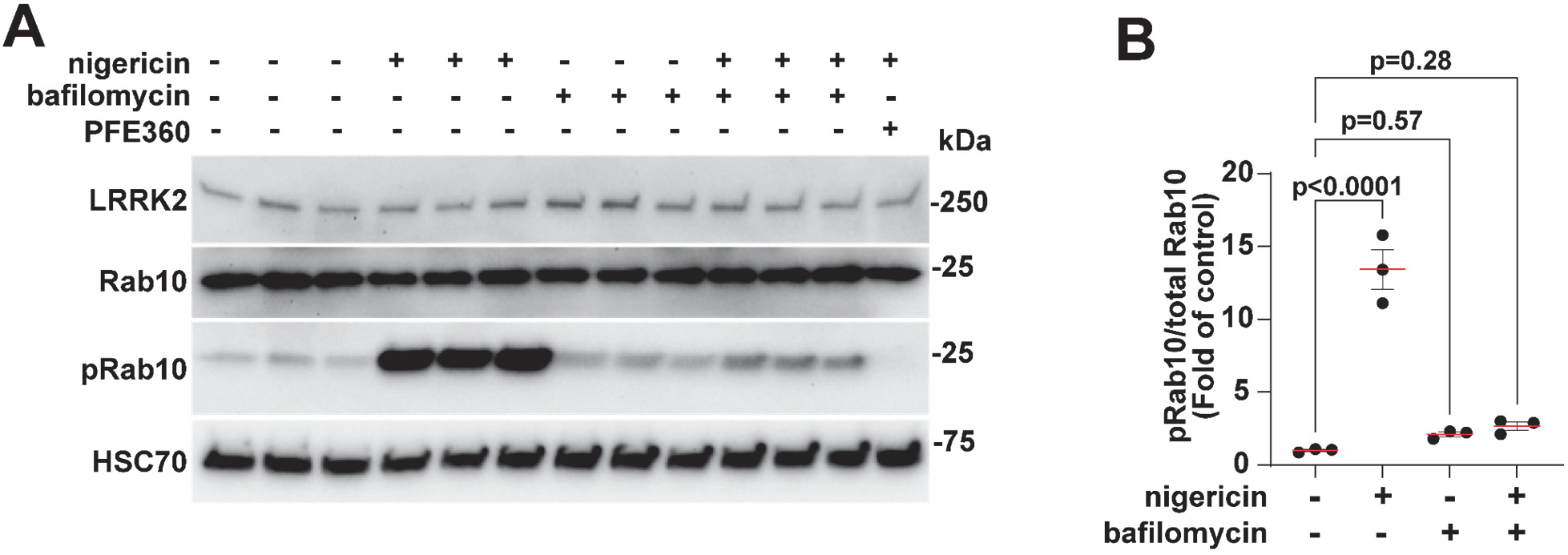
Nigericin and apilimod induce LRRK2-dependent Rab10 phosphorylation to a similar extent. **(A)** Immunoblots of RAW 264.7 cells treated for 2 h with the ionophore nigericin (10 µM), bafilomycin (200 nM), or the LRRK2 inhibitor PFE-360 (200 nM) alone or in combination as indicated. Blots were probed for LRRK2, Rab10, pRab10, and Hsc70 as a loading control. **(B)** Quantification of pRab10/Rab10 ratios. Nigericin strongly increased pRab10, an effect abolished by PFE-360, confirming LRRK2 dependency. Bafilomycin alone or in combination with nigericin had no effect. Each dot represents immunoblot analysis of one biological replicate.

**Figure S5.**
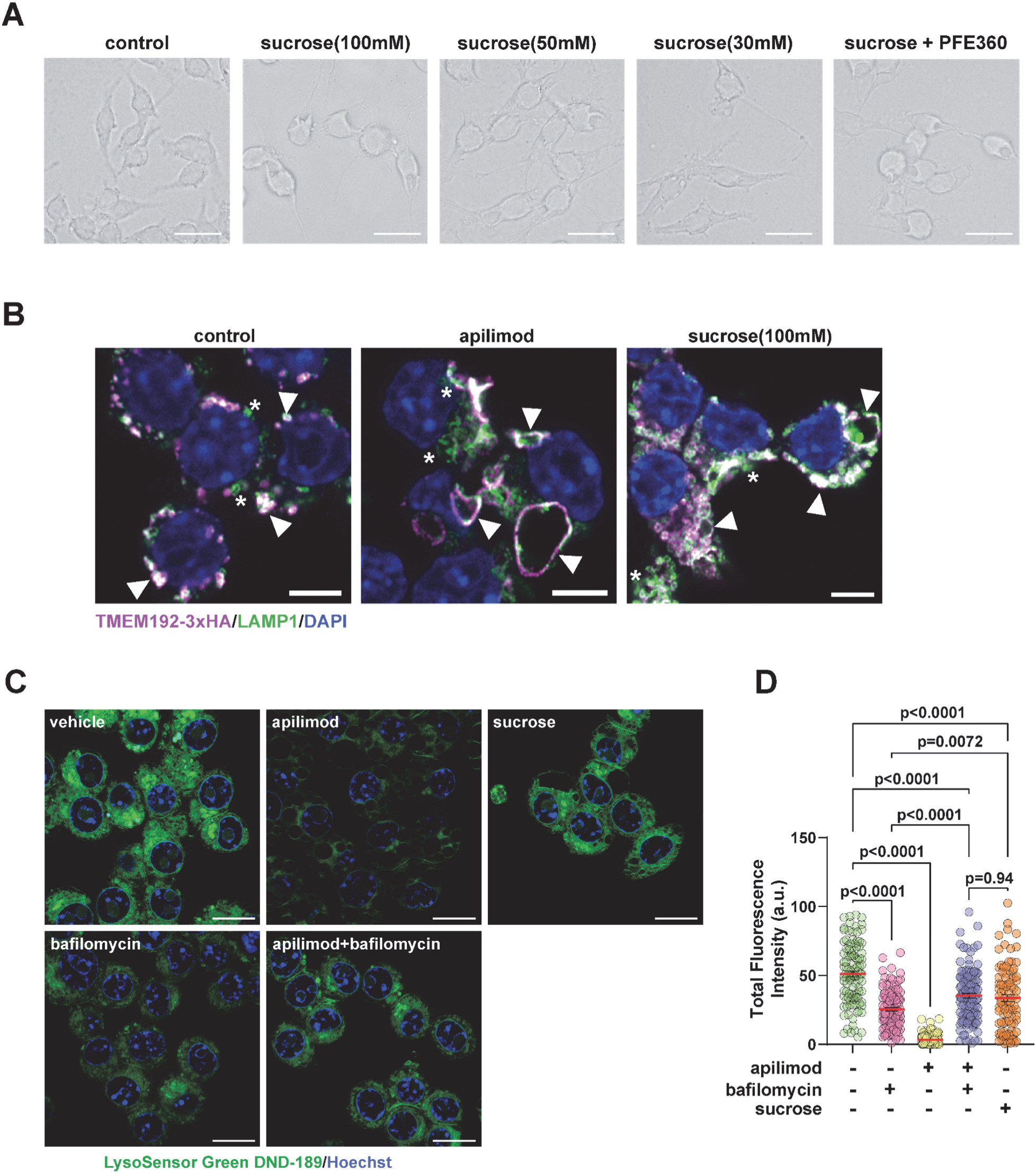
Lysosomes are comparably swollen after both apilimod and sucrose treatments. **(A)** Representative brightfield images of RAW 264.7 cells after 24 h exposure to vehicle (DMSO, 0.01%), sucrose at 100, 50, or 30 mM, and sucrose (100mM)+PFE-360. Scale bar, 20 µm. **(B)** Representative confocal images of RAW 264.7 macrophages expressing TMEM192–3×HA (magenta) and stained for LAMP1 (green) and DAPI (blue) following 2 h treatment with vehicle, apilimod (100 nM), or sucrose (24 h, 100 mM). Asterisks mark LAMP1–positive compartments and arrowheads indicate TMEM192–3×HA-poisitive/LAMP1-positive compartments. Both apilimod and sucrose caused evident lysosomal enlargement and partial separation of TMEM192–3×HA and LAMP1 signals, consistent with swollen or distended lysosomes. Scale bar, 20 µm. **(C)** Representative confocal images of RAW 264.7 macrophages stained with LysoSensor Green DND-189 (green) and Hoechst 33342 (blue) after 2 h treatment with vehicle, apilimod (100 nM), bafilomycin (200 nM), apilimod + bafilomycin, and 24 h treatment with sucrose (100 mM). Apilimod and bafilomycin markedly reduced LysoSensor fluorescence, indicating elevated lysosomal pH, whereas sucrose produced moderate fluorescence loss, consistent with partial neutralization. Scale bar, 10 µm. **(D)** Quantification of total LysoSensor fluorescence intensity per cell. Both apilimod and bafilomycin significantly decreased fluorescence relative to vehicle, while co-treatment did not further reduce intensity. Sucrose caused a smaller but significant reduction. Data are mean ± SEM, *N* = 3 independent biological replicates, one-way ANOVA test. At least 30 cells per one biological replicate of staining analysis were analyzed.

